# Restoration, dispersal and settlement of native European oyster (*Ostrea edulis)* in energetic tidal areas

**DOI:** 10.1101/853911

**Authors:** Jay Willis, Lisa Kamphausen, Anthony Jensen, Roger J.H. Herbert, Elfed Jones

**Affiliations:** Scottish Natural Heritage, Inverness, IV3 8NW UK; University of Southampton, SO17 1BJ UK; University of Bournemouth, BH1 3LT UK; HR Wallingford Ltd. OX10 8BA UK

**Keywords:** Benign invasion, Connectivity, ELAM Eulerian Lagrangian Agent Based Model, Habitat restoration, PLD Pelagic larval duration, Range collapse, Spat fall

## Abstract

We modelled the pelagic larval phase of native European oyster (*Ostrea edulis*) in the Solent. The Solent is a complex and tidally energetic environment on the south coast of the UK. Until recently it was the largest self-sustaining fishery of the native oyster in Europe. We developed a new larval settlement behavioural model that is the simplest plausible model which remains consistent with all available data and evidence on larval behaviour. We used a hydrodynamic sub-model, a Lagrangian advection sub-model, and an individual agent based model. The results demonstrate how isolated oyster assemblages can re-populate larger areas of historical inhabitance. We predict the most likely patterns of redistribution from refugia or from fisheries seeding. We show that settlement swimming behaviour is as equally important as passive hydrodynamic transport for larval survival and adult distribution and that settlement swimming behaviour has a profound impact on settlement patterns. The models show that managed broodstock refugia have the potential to seed much larger oyster beds in contrast to broad-scale seeding. Such refugia have the advantage of maintaining a locally high and mature population with potential for reef features and their associated biodiversity. We show that such refugia may be best placed in the tidally dynamic and exposed areas rather than on sheltered coastal sites as they have been in the past. Our model is insensitive to parameter variation and could be an effective and practical management tool in the face of a paucity of field data on larval distribution and behaviour.

## Introduction

Native oysters (*Ostrea edulis.* Linnaeus, 1758) used to be common along the south coast of England (Gardner & Elliot 2001) and now there are only isolated groups (Laing *et al.* 2005). To restore and conserve fished stocks and native oyster reefs, it is vital to know if the original populations were connected or self sustaining and to understand how their larvae are potentially transported during the dispersal stage of their life history. *Ostrea edulis* have a 10 - 16 day pelagic larval phase (Waller 1981). During this phase they could be transported hundreds of kilometres in the strong tidal currents of the English Channel, as has been shown with other bivalves (Herbert *et al.* 2012). If *Ostrea edulis* on the south coast are dependent on larval supply from each other, i.e. the oysters are a meta-population (Kritzer & Sale 2004), restoration attempts of isolated populations are only likely to be temporary solutions.

We have aimed for a model of behaviour which is consistent with all known scientific study, and which is plausible; by which we mean that we have not assumed any knowledge, memory, physiological or sensory capability that has not been demonstrated for this species (Andrews 1979, Waller 1981, Cole & Knight-Jones 1939 and 1949, Korringa 1941, Bayne 1969, Walne 1974, Andrews 1979, Galtsoff 1964, Cragg & Gruffydd 1975). *Ostrea edulis* larvae are strong and consistent swimmers but little is known about their cognitive or sensory capabilities and larvae of different but closely related aquatic species often exhibit quite different swimming behaviour (Leis 2006). We tried to avoid providing the model larvae with information existent in the model that, in reality, would be unavailable. We avoided models that implied cognitive abilities in the larvae, that had not been observed for this species, but which are undoubtedly present in other navigating animals (Walker *et al.* 2002).

The settlement model incorporated the facts:

1. that young larvae swim almost continuously (milling) but either are evenly distributed in the water column, or target layers of oxygenated water, and in any case staying away from the surface and bed (Cragg & Gruffyd 1975, Waller 1981, Key 1987),
2. settlement aged larvae settled on the bed in the default laboratory conditions (Cragg & Gruffyd 1975),
3. settlement aged larvae in the lab responded to very small pressure stimuli (equivalent to less than 10 cm water depth change) by swimming up off the bed in increasing pressure and returning back after pressure is reduced (Cragg & Gruffyd 1975), and
4. younger smaller larvae are likely to be eaten by adults of the same species (Troost *et al.* 2008).

These led to the minimum pelagic swimming and settlement model, which consisted of neutral buoyancy and a random walk swimming model in the pelagic phase (which causes transport away from natal release zone and adults and which is important in respect of point 4 above) followed by trial settlement and bouncing, when depth increases and larvae are on the bed, in the settlement stage (which leads to movement toward suitable settlement habitat (on average)). This model, as a hypothesis, had the benefit of being consistent with recorded observations (Cranfield 1973, Cragg & Gruffyd 1975, Waller 1981, Key 1987, Cole & Knight-Jones 1949, Korringa 1941, Bayne 1969, Walne 1974, Andrews 1979, Galtsoff 1964) and conversely is not falsified by them, and of being plausible, in that it has been shown that larvae would and could respond in this way. Furthermore, it does not incorporate abilities that have been shown for other species (such as endogenous timing of tides or response to salinity gradients) which would be inconsistent with observations of this species as evenly distributed through the water column in all tide states (Key 1987).

In the Solent all this is of particular interest. Native oysters have been cultivated in the Solent since Roman times (Günther 1897). Solent oyster populations have frequently collapsed and recovered (e.g. Orton 1923, Orton 1927, Tubbs 1999). The most recent collapse happened in 1962 – 63. Beds in the Beaulieu river were re-stocked, and it is assumed that successive good spatfalls from this source triggered a recovery in the Solent and established the current population. In 1972 approximately 2 million (110 tons) of oysters over 38 mm were caught by commercial fishing (Davidson 1976). Throughout the 1970s and 80s the population of *Ostrea edulis* recovered further and spread through the Solent which became the largest so-called natural *Ostrea edulis* fishery in Europe (Tubbs 1999). It peaked in the 1979-80 season, with landings in Solent ports of about 840 tonnes, or 15 million oysters (Key & Davidson, 1981). Since then catches have been falling considerably every year. The oyster beds in the Beaulieu River were cleared out by the government fisheries authorities in 1986 as a bonamiosis control measure. In 2010 landings data from the UK Marine Management Organisation sea fisheries statistics recorded 87.57 tonnes of oysters landed in Portsmouth. Some additional landings will have been made in other places, but unfortunately no comprehensive data on exact catches from the Solent is available. The Solent oyster fishery order expired in 2010, and in the light of a rapidly contracting stock, regulatory authorities and stakeholders agreed not to renew it.

An annual stock survey has been conducted since the 1970s. Catch rates were highest along the western mainland shore of the Solent during the peak in the late 1970s, but catches gradually moved into the eastern Solent, until from about 2000 onwards all commercially fishable grounds were located there (Vanstaen & Palmer 2009). Today the beds at Sowley, Lepe and Lymington in the West are effectively barren, and Ryde Middle, Warner and Browndown in the East yield the largest remaining catches (Vanstaen & Palmer, 2010). Reasons for this perceived shift of the stock from the West Solent into the East Solent are unclear, but suggestions have been made ranging from pollutants being introduced into the West Solent, to prevailing currents carrying bacteria from decomposing red tides in Southampton Water into the West Solent and inhibiting recruitment (Crawford *et al.* 1993).

The catastrophic long term loss of areal extent and biomass of oyster grounds are global challenges. Studies using fisheries data almost universally fail to quantify loss, and tend to use baselines on populations which have been already subject to long term exploitation (Ermgasson *et al.* 2012). Biomass declines are greater than areal extent and both are usually very far from the pristine condition (Ermgasson *et al.* 2013). Thus relatively higher abundances in the Solent, noted above, may be far from the true pristine carrying capacity and establishment of reefs with longer lived individuals may serve to buffer against the existing variability. Reefs may also provide refuge from storm surges and high wave events. There may be significant unknown benefits of much higher biomass aggregations, to both the oyster population and biodiversity. Biogenic oyster reefs which armour the bed, change local sedimentary and water transport patterns and provide habitat structure for a range of species which may presently have none in this area (Ermgasson *et al.* 2013). In short we should be prepared to aim for conservation targets in terms of biomass (and not areal extent) well above what might ever have been documented in the past.

## Methods

### Study area

Between the Isle of Wight and mainland shore (Fig. 1) the Solent receives fluvial discharge from several rivers that rise in farmland and flow through both small recreational and large industrial harbours. There are strong west-east tidal streams (max 3ms-1) that cause considerable mixing. Mean salinity is 34 and sea temperature range is 6-19°C. Sediments on the oyster beds vary and consist of muds and coarser grained sand and gravels.

**Figure 1.**
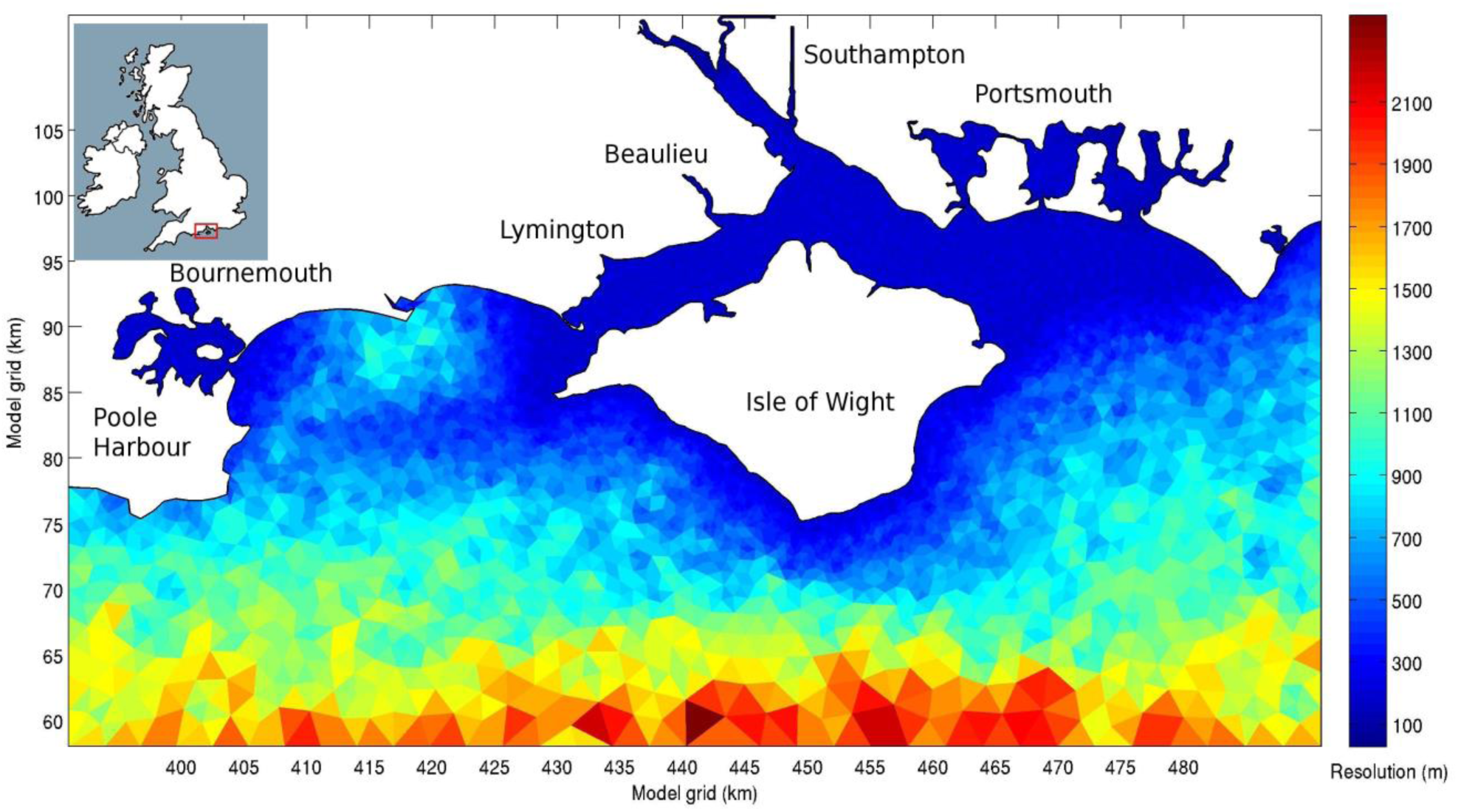
The Solent on the south coast of the UK with boundary at limits of the hydrodynamic model (red box on inset). Spatial resolution is shown for each hydrodynamic model triangular cell. The resolution out in the channel (to the south) is several km, whereas in the Solent (between the Isle of Wight and the mainland) it is less than 100 m.

### Water model

Hydrodynamic conditions in the study area (Fig. 1) were simulated using the two-dimensional (2D), depth-averaged flow model TELEMAC-2D (http://www.opentelemac.org/). The 2D hydrodynamic model was built and calibrated using a variety of existing and newly surveyed bathymetric and flow data. The spatial resolution of the unstructured triangular mesh used by the model in the Solent is variable between about 100 m and 2000 m. The water model was run ‘off-line’ for a full spring-neap tidal cycle and the velocity and depth results at 1400 s time steps were saved. A start time (after the model had stabilised (run-in)) and an end time were chosen when the tide had returned as close as possible to a similar state, including the full spring-neap tidal cycle, and this information was used to ‘loop’ the water model indefinitely in time.

### Passive drifting Lagrangian particle model

The methods used for modelling swimming animals and deriving a quasi-3D model for particle dispersion from a 2D hydrodynamic model are outlined in Herbert *et al.* (2012) and follow Mead (2004, 2008) and the recommendations of the relevant ICES working group (North *et al.* 2009, Willis 2011). We used Runge Kutta fourth order integration to calculate the velocity of the particles from the water velocities (Willis 2011).

### Initialisation

For each scenario 2500 larvae were released simultaneously from a set of release areas (Fig. 2), multiple runs of 10 × 2500 particles were used where key results of a single run were numerically low to give better resolution of uncertainty (North *et al. 2009).* Release areas were either ‘beds’ of known habitation or square experimental ‘zones’ (related to field studies). Positions of oyster beds were digitised into the model domain using the 2011 report (Palmer & Firmin, 2011). Initial positions within the release areas were randomly chosen. The model larvae remained in the model for the pelagic phase and the settlement phase (14 days in total). Their positions every hour were recorded. Timing of release (in respect of tides) was spread randomly over a release period (default 4 hours). All model larvae were assigned an initial size of 160 *μ*m and a growth rate. The growth rates were randomly picked from a normal random distribution with a mean that ensured, on average, that the model larvae reach 300 *μ*m after 10 days, and with a variance of 1 day. Once each larva had reached 300 *μ*m it switched its behaviour from pelagic swimming (a random walk based on the milling speeds at age reported by Cragg and Gruffyd (1975) and neutral buoyancy) to settlement behaviour. This size model was used to provide a realistic dispersion around the time to settlement and for calculating swim speed. In the future it could be moderated by the modelled fate of the individual larvae so that time spent at certain depths, salinities or temperatures could impact the growth of individuals.

**Figure 2.**
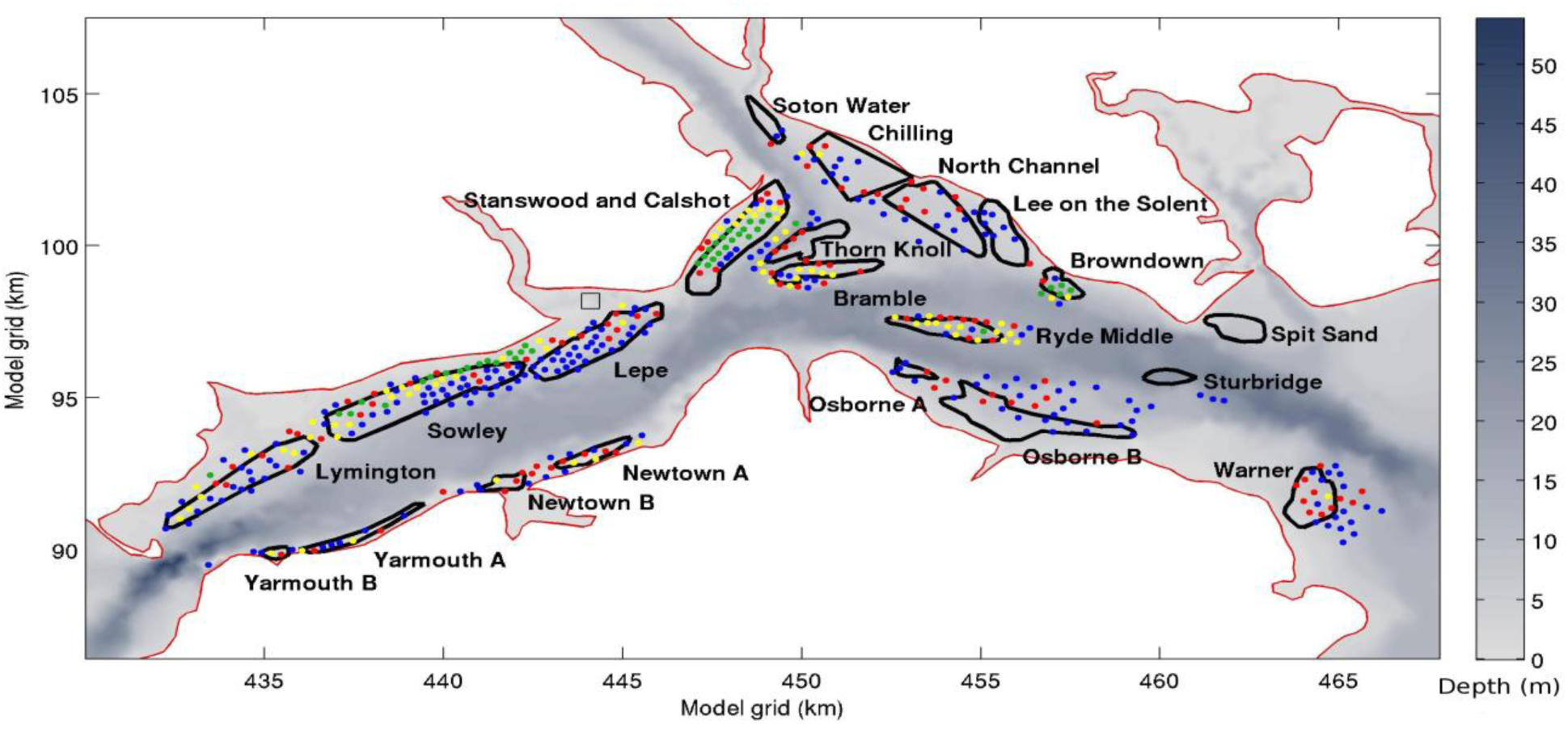
Oyster distribution areas and release zones in the Solent. The named beds are digitised from Palmer and Firmin (2011) and the coloured dots from an earlier dredge survey when the Solent oyster population was at a peak (Key & Davidson, 1981), where blue = 1-5, yellow = 6-14, green = 15-49, red = >50 oysters. The small square zone above the Lepe bed is the Beaulieu refuge start zone.

### Settlement model

The settlement state was initialised by swimming down to the bed. From then on, until the larvae settled, they ‘bounced up’ from the bed during times of increasing water pressure, i.e. they swam upward continually for a pre-set time after they had sensed that the depth was increasing (swim speed range at length derived from: Hidu & Haskin 1978, Cragg & Gruffyd 1975). After the bounce time had elapsed the larvae swam downward, back to the bed. Once returned to the bed they bounced again if the depth was still increasing, and not if depth was stable or decreasing. Literature on *Ostrea edulis* larval behaviour supports these assumptions (Cranfield 1973, Cragg & Gruffyd 1975, Waller 1981, Key 1987). The second element of the settlement model involves testing the bed. The suitability of the present location for settlement was incorporated in the model as a probability of settlement assigned to every location. Each time a model larva was near the bed it tested the settlement surface nearest to its location with a uniform random number generator, if the random generated number was within the settlement range the larva settled and moved no further in the model. A multiplier was used to ensure that settlement of all the model larvae was roughly spread over the settlement period known for this species (4 days, e.g. Walne 1964 and 1974, Key 1987). The model was predictably and linearly sensitive to this multiplier, which, after testing, was fixed to allow comparison between treatments.

### Settlement surface

The settlement surface was built on a 100 m square grid over the modelled area, using the same underlying OSGB grid as the hydrodynamic model (Fig. 4). The settlement surface was determined by the historical distribution of the species at the time of the last population peak. Dredge data collected in 1981 (Key & Davidson 1981) were digitised from sketch maps and were manually positioned in the model domain onto the 100 m grid of the settlement surface using a standard image manipulation program (GIMP http://www.gimp.org/). Catch counts had been recorded in ranges (0-5, 6-10, and so forth) and the median value was placed within the corresponding grid cell. The positions of known oyster beds from before the current population collapsed were similarly digitised into the model domain using the 2011 report (Palmer & Firmin 2011). These beds were also used as initialisation zones in certain model scenarios. Settlement surface grid cells which were within any of the known beds were assigned an equivalent dredge catch count of 1. A Gaussian spreader of extent 300 m (standard deviation about 50 m) was then applied to all the point data to create a smoothed surface on the settlement grid, which was normalised to a maximum height for any cell of 1 (the minimum was zero).

**Figure 3.**
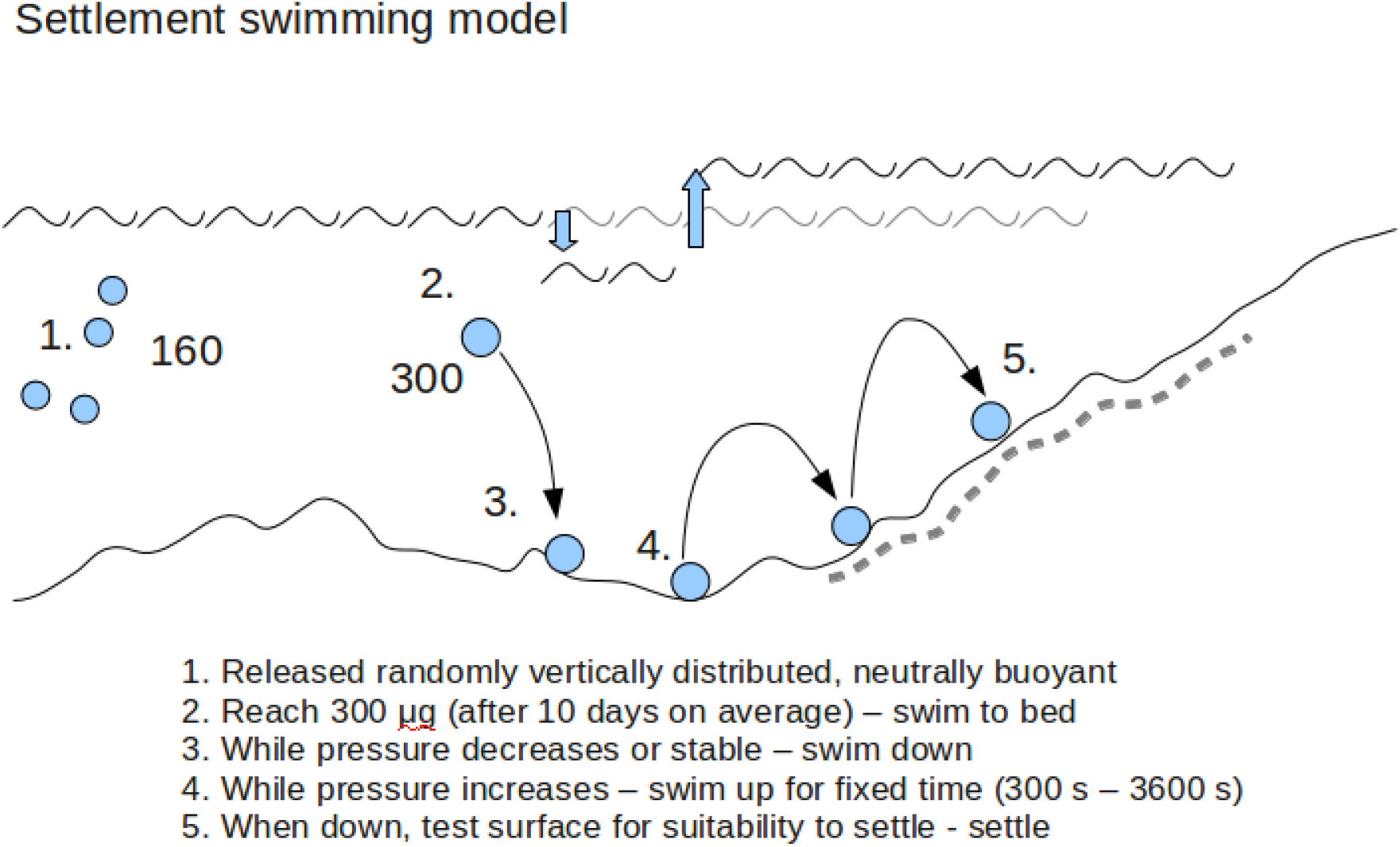
Diagrammatic representation of the settlement swimming model. At first the particles (oyster larvae) are neutrally buoyant and distribute randomly in the water column. They start at a weight of 160 micrograms on average (situation 1.) (Each has a different starting weight chosen from a normal random distribution). They grow as the model progresses and when they reach the threshold on 300 micrograms (situation 2.), they swim to the bed. Thence onward, they swim down when the pressure decreases or is stable – this is when the tide is slack or going out, and when the pressure is increasing (tide coming in) they bounce up into the water column, swimming upward for a variable amount of time, and then downward again after the time elapes. Each time they reach the bed, they ‘test’ the bed – a random chance of settlement is calculated based on the settlement surface. This behaviour matches laboratory responses of larvae to pressure changes and clearly is expected to have the effect of moving larvae to shallower water as they will moved by the currents only during the bounce.

**Figure 4.**
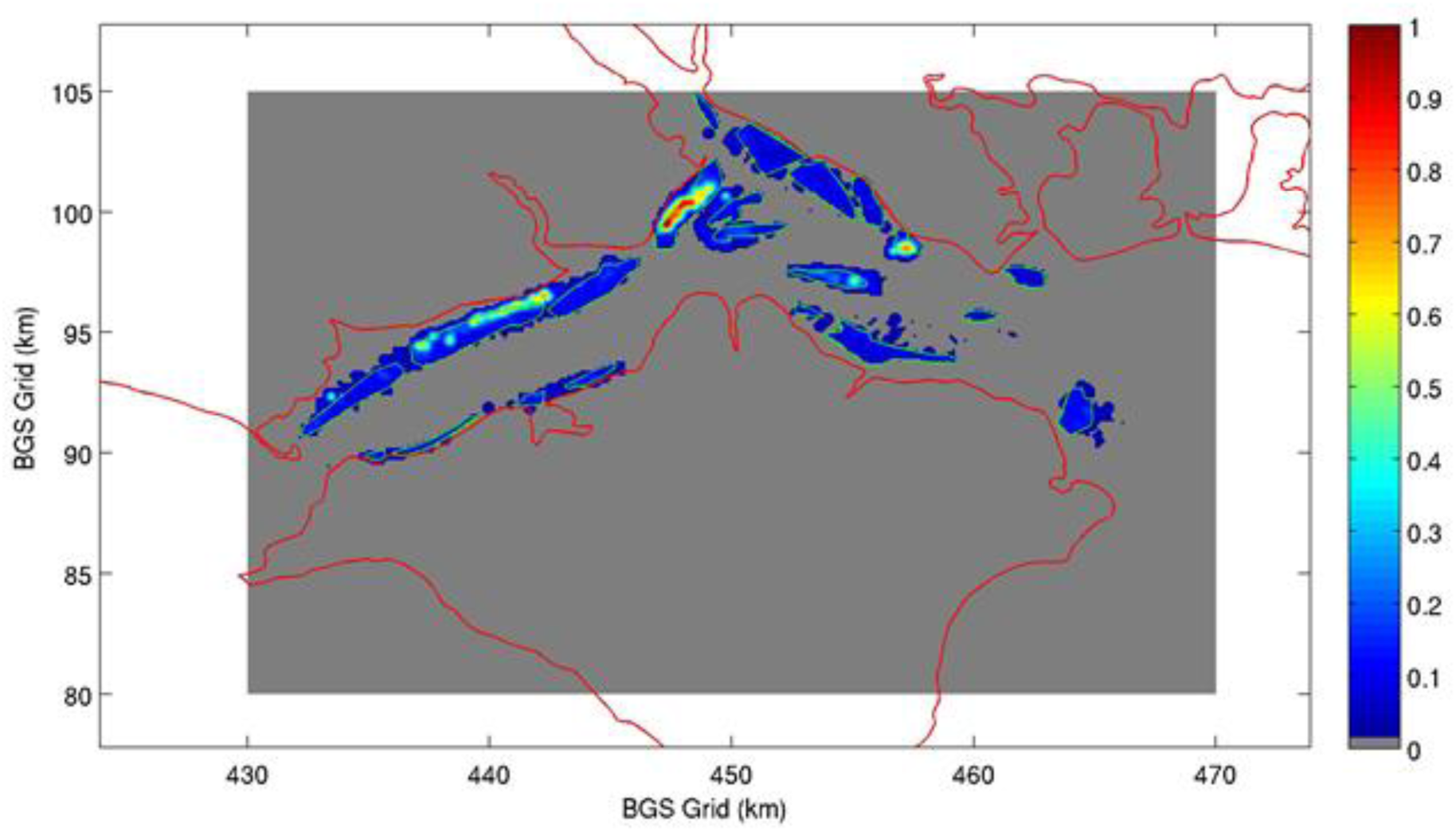
Settlement surface. The value on the settlement surface lies between 0 and 1 and represents the probability of settlement if the model larva ‘tests’ the bed within that grid square. The surface is on a 100 m square grid and is created by addition of the named beds and dredge surveys (Figure 2). The beds contribute a uniform value and the dredge survey results, which are point data, are made into a surface through the use of a Gaussian spreader of 300 m in extent. The grey box represents the extent of the settlement surface – outside this area no settlement occurred (in the model).

### Scenarios

The scenarios tested in this modelling were:

1. Release from specific refuge zone (Beaulieu), without settlement model.
2. Release from beds of previous habitation, without settlement model.
3. Release from beds of previous habitation, with settlement model.
4. Release from beds of previous habitation with long bounce settlement model.

The first scenario was designed to test the patterns of dispersion that might arise from small isolated refuge populations and contrast these with the patterns, developed in scenario 2, which emerge from a population in as close to a pristine distribution as we can estimate. The settlement model scenarios were designed to test if the settlement model was effective in terms of alteration of the patterns of dispersion and if so, to what extent these might improve survival likelihood (settlement in existing areas of habitation) of larvae.

### Parameters

The sub-models each have a wide variety of variable parameters. For the majority of physical parameters of the water model including the Lagrangian drifting sub-model we used parameters that had been calibrated or proven effective in past studies which use the same techniques and modelling software (Herbert *et al.* 2012, Willis 2012). In the case of parameters related to the oyster larvae, which were unknown, such as precise location of spawning, tidal state at spawning or length of pelagic swimming phase, we used a range of values randomly selected around a baseline parameter set.

### Statistics

The variation in the probability of settlement was calculated by randomly re-sampling 10 groups of 500 from the 2500 independent results of each model run. The time to settle statistic more closely followed a gamma distribution than a normal, as determined by maximum likelihood. We give the mean and standard deviation derived from the best fit gamma distribution in these cases. For the passive drifting cases we calculated probability of settlement by counting all the hourly recorded positions of 2500 particles, during 3 day settlement period (180,000 positions), that were over settlement surface (any grid square greater than zero on the settlement surface) and divided the number by all the positions. The mean and variances were calculated by re-sampling 10 groups of 500 particles as above.

### Sensitivity

The key model result (settlement probability) was tested for sensitivity by running multiple full scenarios with variation of relevant parameters (coefficient of diffusion, vertical diffusion, tidal state at start, milling speed, range of swimming speed, range of settlement times, step frequency for settlement testing, and tidal sensitivity). Each model run was analysed in the standard way by re-sampling 10 groups and recording mean and standard deviation. The sensitivity run statistics were then compared to each other and to the default run to identify any potential significant differences, using a two sample t-test or observation of the standard deviations.

## Results

The larvae moved on average 322 km before settlement (standard deviation 140 km, N=2500). The probability of settlement was significantly higher for bounce models than for passive drifting (Table 1). Between the long (30 minute) and the short (5 minute) bounce models there was no significant difference. The probability of settlement when larvae were released from the refuge area in the Beaulieu, using passive drifting, was very low (3.2%) but significantly higher than from the assumed pristine distribution. The distances travelled by settlers between points of initialisation and settlement were well modelled by a gamma distribution (chosen using maximum likelihood) with mean of 9.7 km, standard deviation 7.6 km, range 1-30 km..

**Table 1.** Results for each of the scenarios, mean with standard deviations in parenthesis, for 10 re-sampled groups of 500 from 2500.

| Scenario | Probability of settlement (%) | Mean time to settlement (hours) | Mean depth at settlement (m) |
| --- | --- | --- | --- |
| 1. Passive from refuge | 3.20 (0.09) | – | – |
| 2. Passive from pristine | 1.33 (0.13) | – | – |
| 3. Bounce settlement | 40.64 (1.9) | 24 (25) | 13.01 (5.4) |
| 4. Long bounce | 37.62 (2.9) | 22 (24) | 11.91 (5.7) |

### Dispersion kernels and settlement patterns

Dispersion kernels were constructed for the passive drifting scenarios, without a settlement model, by recording hourly positions throughout the final 3 days of the model run, this led to 180,000 positions which were then counted on a grid across the central modelled area, linearly interpolated, and normalised such that the entire volume of the resultant surface summed to unity, to make this a probability density kernel. The logarithm of this surface tends to accentuate the less dense areas to produce a smoothed pattern over the area which is informative regarding the general dispersion (Fig. 5, Fig. 6).

**Figure 5.**
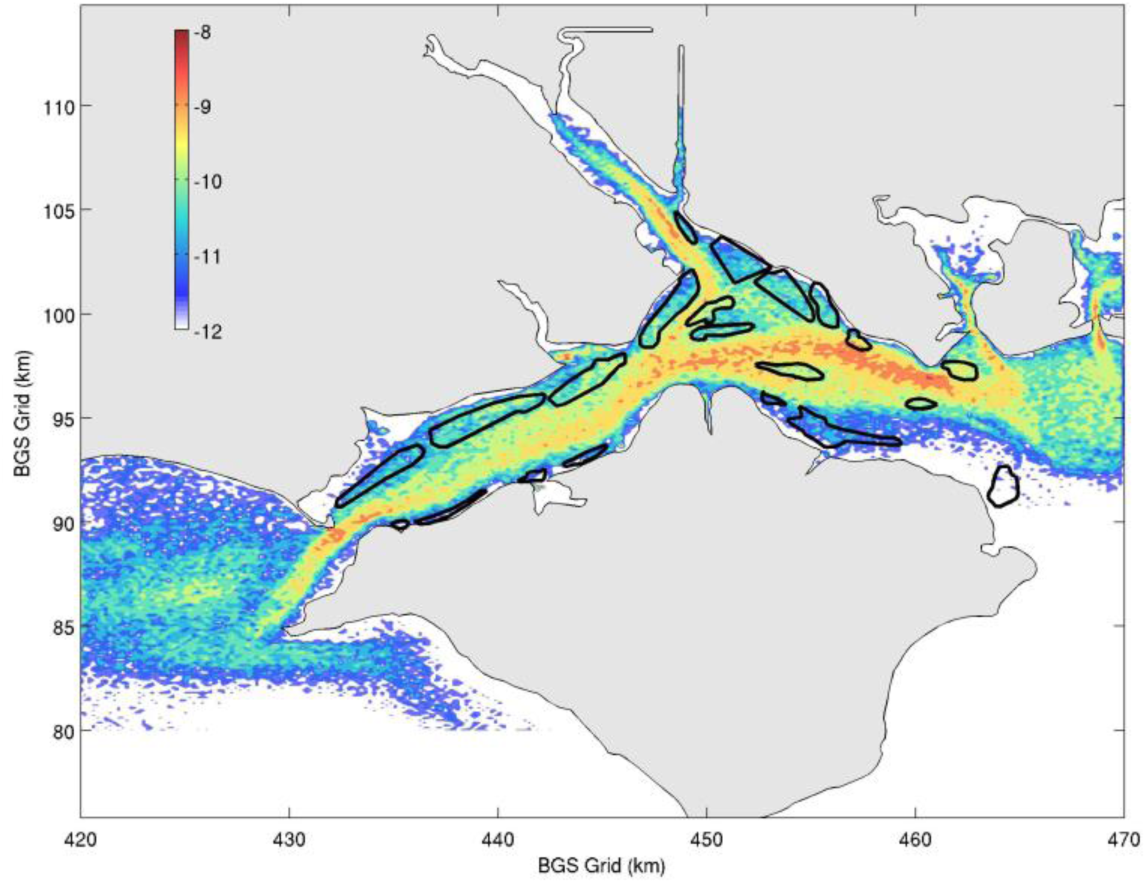
Log transformed kernel density estimate (coloured surface) of all hourly positions in the last three days of a 14 day passive drifting advection model of 2500 particles (N=180,000). The log transformation accentuates the areas of lower density. The particles were initially released from the beds outlined in thick black lines. The density of the particles is well correlated to bathymetry in the Solent (Fig. 2), and there are relatively few positions recorded over the beds, where larvae are expected to settle.

**Figure 6.**
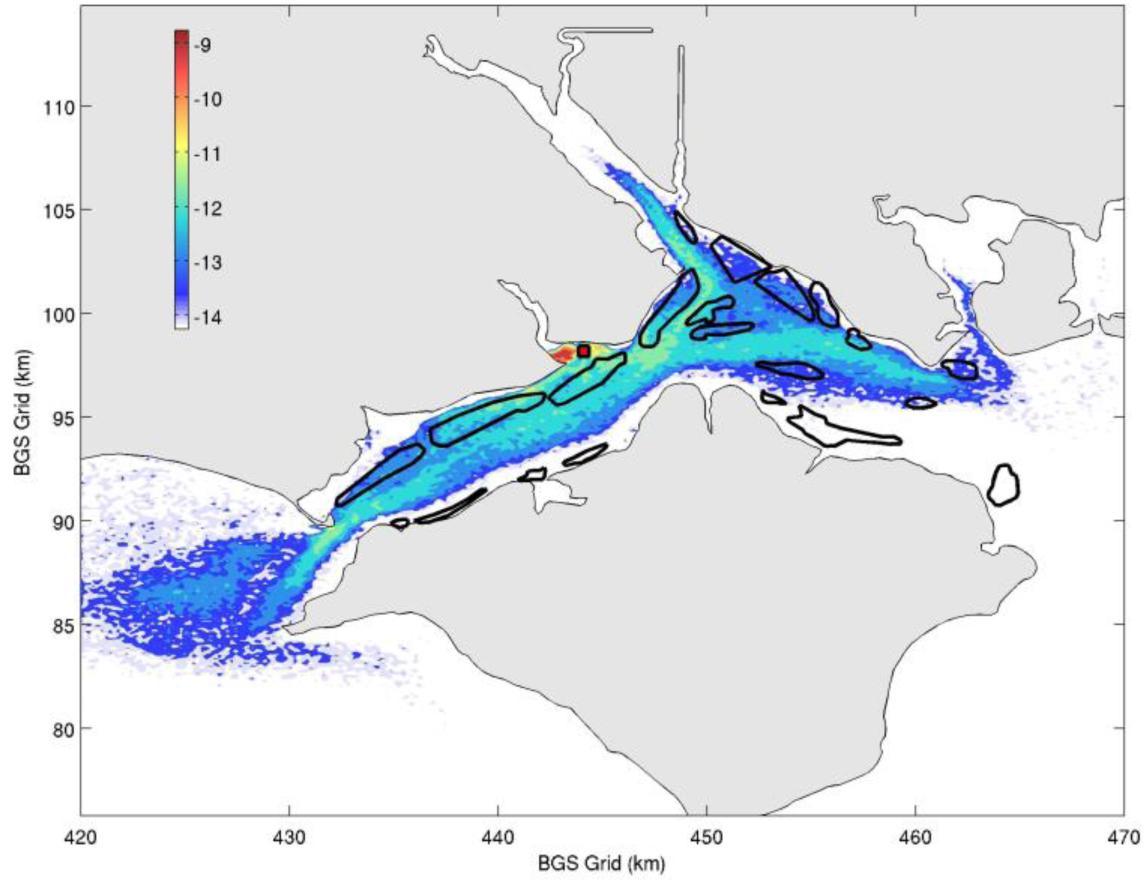
Log transformed kernel density estimate (coloured surface) of all hourly positions in the last three days of a 14 day passive drifting advection model of 2500 particles (N=180,000). The log transformation accentuates the areas of lower density. The particles were initially released from the 500 m square zone (red square) just on the exit of the Beaulieu estuary. The density of the particles that left the estuary is well correlated to bathymetry in the Solent (Fig. 2), and there are relatively few positions recorded over the beds, where larvae are expected to settle.

### Settlement model performance

A typical dispersion kernel representation of settlement positions after settlement model shows well distributed settlement in all the beds (Fig. 7). The ratio of initialisation numbers to settlement numbers for each bed highlights those beds that attracted a higher density of settlement independently of their areal extent (Table 2). There was a hot spot in and near Browndown (Fig. 7). All beds imported more larvae than they exported (Table 2), usually 5-15 times more and only Ryde Middle was both a high relative self seeder and importer. The other high importers were relatively low self seeders. Individual journeys for model larvae were represented either in the vertical dimension (Fig. 8 left panel), or they can be represent plan view, where the x and y coordinates recorded at each output step for each larvae show their horizontal passage through the area (Fig. 8 right panel). Settlement journeys were very varied (Fig. 9). The larvae were randomly and evenly distributed in the water column before they reached settlement, and each larva moved through the whole water column. Many larvae left the Solent and re-entered during the settlement phase.

**Table 2.** Initialisation to settlement ratios using the 1800 s long bounce settlement model (n=10×25,000). Initial density was equal across the beds but they are different sizes so proportions of numerical values are more informative about the beds relative attraction. Import indicates all settlement ratio to initialisation including self seeded larvae (larvae which both initialised and settled in the same bed). Many model larvae settled outside of named beds (Fig. 2) (especially near to Browndown). Bold names show initialisation to settlement ratio over 0.5, and proportion of self settlement over 0.05.

| Bed name | Import | Self seed (x 10<sup>-2</sup>) |
| --- | --- | --- |
| Lymington | 0.11 | 2.1 |
| Sowley | 0.18 | **5.2** |
| Lepe | 0.14 | 2.9 |
| Stanswood and Calshot | 0.11 | 2.2 |
| Soton Water | 0.15 | 0 |
| Chilling | 0.03 | 0.3 |
| North Channel | 0.12 | 0.4 |
| Thorn Knoll | 0.3 | 0.9 |
| Bramble | 0.17 | 0.8 |
| **Ryde Middle** | **0.63** | **6.6** |
| Lee on the Solent | 0.07 | 0.1 |
| **Browndown** | **0.82** | 0 |
| **Spit Sand** | **0.53** | 0 |
| **Sturbridge** | **0.73** | 1.5 |
| Warner | 0.02 | 1.4 |
| Osborne A | 0.3 | 0.4 |
| Osborne B | 0.27 | **6.4** |
| Newtown A | 0.14 | **6.1** |
| Newtown B | 0.07 | 0.7 |
| Yarmouth A | 0.01 | 1.9 |
| Yarmouth B | 0.01 | 0.7 |

**Figure 7.**
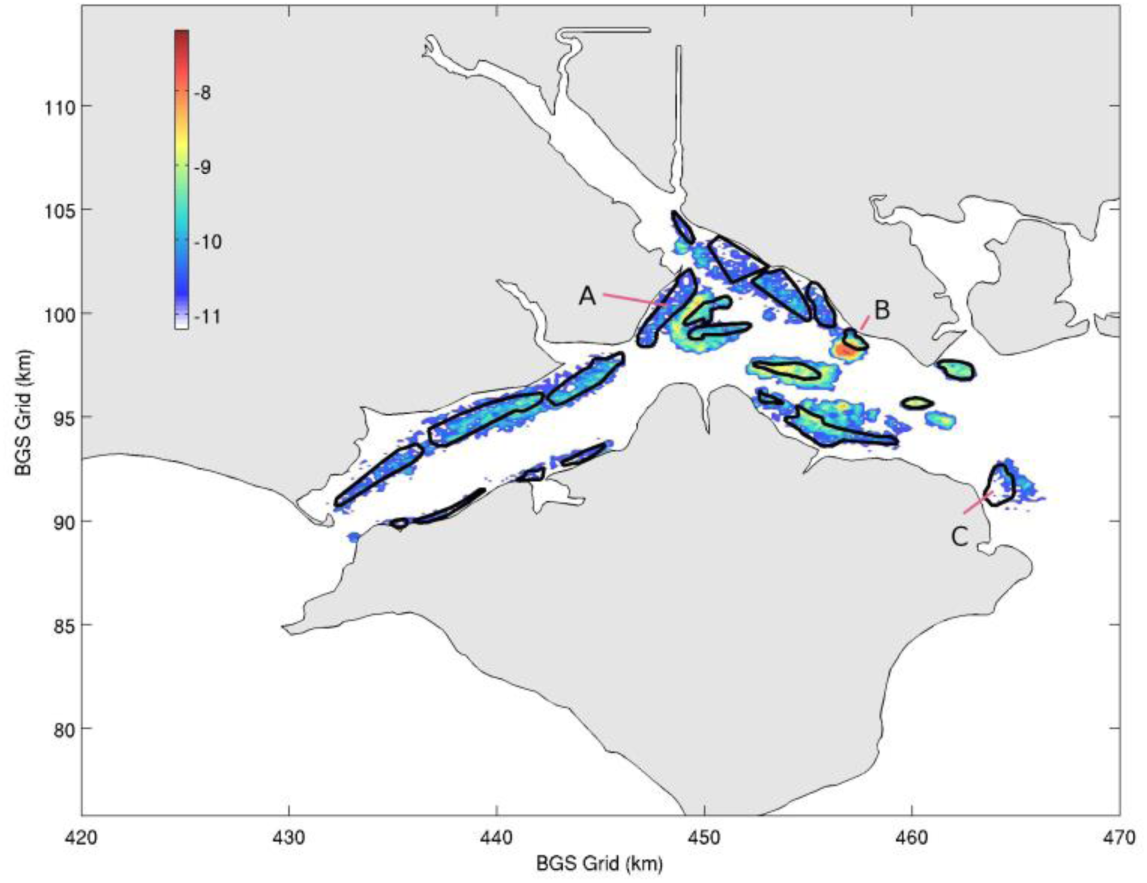
Log transformed dispersion kernel of the settlement probability of model larvae (N=10,289) from original cohort of 25,000 (10 standard scenarios) using the 1800 s, long bounce, settlement model. Particles were initiated randomly placed in the beds (thick black outlined zones). The overall pattern is well distributed settlement across all areas which are known as oyster beds in the Solent. Settlement is less dense in Stanswood and Calshott (A) than would be expected from the observed historic distribution, but the higher survey observations there are probably due to relaying of oysters in the several order fisheries there. There is a strong hot spot at Browndown Bank (B), and generally high settlement on the mid-channel beds, with low density in Warner (C).

**Figure 8.**
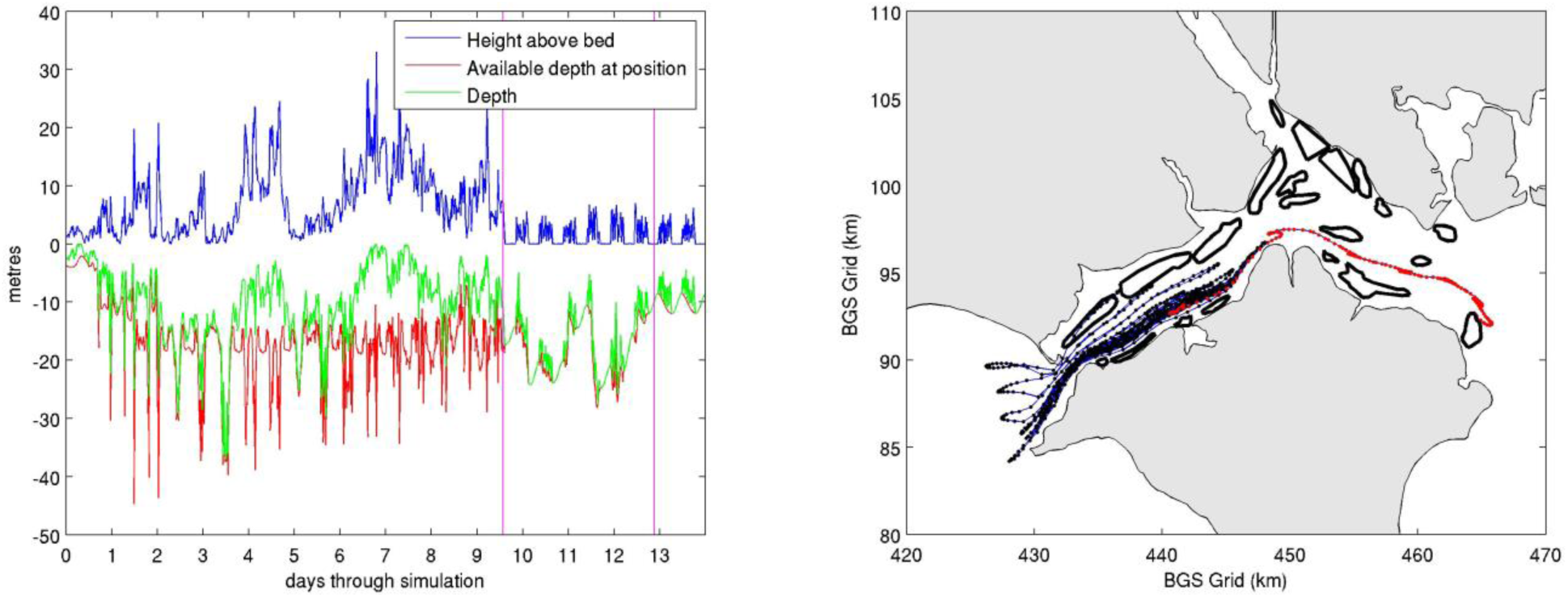
Typical (long) settlement journey for an individual model larva, using the 3600 s long bounce settlement model. Right panel black dots show hourly positions before settlement behaviour, red during. After reaching the settlement size (left panel, left pink vertical line, right panel, left end of red dotted segment) the larva attempts to settle. In the left panel it is evident that around day 11 the larva is bouncing into shallower water, before moving suddenly into deeper water and again moves shallower through day 12 before settlement at the right hand pink line. The right panel shows how the larva moved around 25 km, during settlement behaviour before settling on Warner bank.

**Figure 9.**
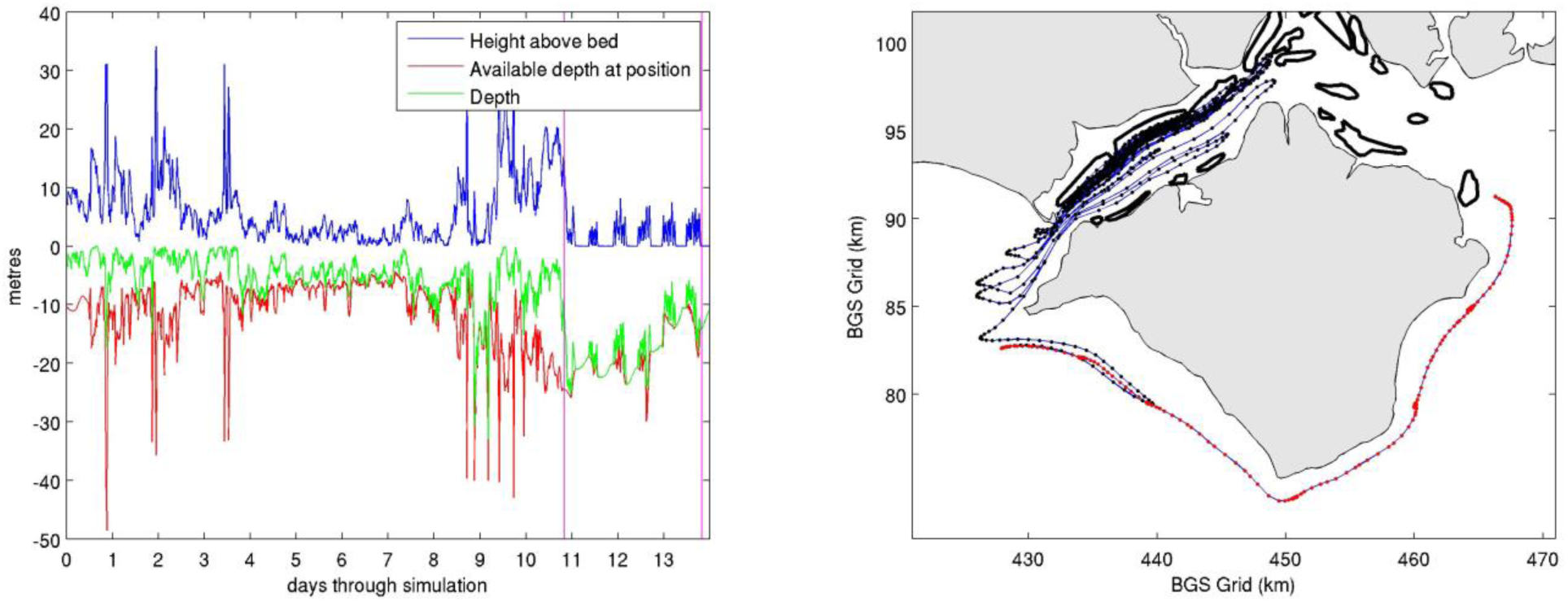
Atypical settlement journey, and a very rare circumnavigation of the Isle of Wight. Right panel black dots show hourly positions before settlement behaviour, red during. It was rare in any of the scenarios for particles to leave the Solent from one end and return the other. The left panel shows that throughout the settlement period (between the pick vertical lines) the larva gradually found shallower water before settlement after 3 days.

## Sensitivity Analysis

The model was insensitive to alteration of all but one tested variable parameters. The model was sensitive to length of pelagic phase before settlement – the sooner the onset of settlement the higher the probability of settlement, in rough proportionality, as would be expected. The settlement behavioural model was similarly insensitive to change of the key parameters of swimming speed and bounce time. Swimming speed is only important in relation to how quick larvae get up out of the boundary layer at the bounce start, or back into the boundary layer at the end of the bounce; these two factors tend to compensate for each other. The results were similarly insensitive to time of bounce over a range between 5 minutes to 30 minutes. This was because the incoming tide, the time of increasing pressure, would last for around 4 hours, and so once a bounce was over, a new one would start, and the difference between the bounce times would relate only to the relatively small time, during bouncing, spent in the boundary layer. The final parameter that was subject to sensitivity analysis was the threshold of sensitivity to incoming tide and although we had good experimental evidence for the upper limit of this value (most insensitive level)(Cragg & Gruffydd, 1975) we found that the results were insensitive to the alteration of this parameter as it only served to delay each bounce by a few minutes.

## Discussion

### Model performance

The bounce settlement behaviour worked remarkably well in steering the larvae to suitable areas. The model larvae moved from and to all known beds in patterns that are consistent with the evidence of past inhabitance, suggesting that we have broadly captured a majority of the essential functionality of the real system. This was not the case for a model that only included passive advection where there are known beds that are not crossed by settlement stage larvae from any known areas of inhabitation. The model is plausible in that if larvae benefited by being able to sense the incoming tide, in a highly complex tidal area, by using the pressure stimuli of increasing depth, they would need a fixed baseline pressure. So moving to the bed provides a baseline, and it also provides an opportunity to test the bed for potential settlement. The model unequivocally and positively answers the question of whether this would be a viable method of moving from unsuitable places for settlement, to which larvae may have drifted, back to suitable areas, within the times of known pelagic behaviour. The striking aspect of the study was the insensitivity of the model to changes in the variable parameters. Since many of the true analogues to the model parameters are unknown this insensitivity is especially valuable as an indication of the model’s likely good predictive capability. Here following we discuss some of the implications of the results assuming they are broadly correct with respect to the actual situation and make some suggestions for future research that could support or directly falsify our hypothesis. Nevertheless we should always caution that there may be things which we have not modelled that could fundamentally change or reverse our predictions, or things that we have modelled incorrectly that could do the same.

### Management implications

The models show that oyster larvae released in the Solent generally settle in the Solent at the end of the pelagic and settlement phase, i.e. that the Solent oyster population is not dependent on external larval supply. Passive drifting patterns of larvae released in a single location, e.g. in the Beaulieu Estuary, are similar to those from the widespread release scenarios. This indicates that it is likely that small managed populations (for instance in the Beaulieu) have been able to re-establish to some extent the areal extent of the natural populations, albeit at likely smaller abundances than would have been maintained by original widespread spawning, and perhaps taking two spawning seasons to reach the outlying areas such as Warners from Beaulieu. Thus the model results support the generally held assumption that the present Solent oyster population has been seeded by managed oyster beds in the Beaulieu. It follows that exposed areas where higher concentrations of oysters are maintained (e.g. Ryde Middle) will deliver spat across the Solent. The levels of self-seeding are low for all areas, with settlers arriving from other beds around 5-15 times more numerous than those that emerged locally. The models suggest that it is currently unlikely that natural refugia exist, and as such the Solent oysters could be viewed as a single population across the modelled area. This conclusion is supported by the recent widespread changes in abundance regardless of localized cessation of fishing (e.g. in the Stanswood Bay and Calshot several order fisheries).

### Conservation

New UK legislation under the Marine and Coastal Access Act (2009) makes provisions that allow for spatial protection of *Ostrea edulis* (which is a Biodiversity Action Plan Priority Species (Gardner & Elliot, 2001)). In the Solent, creation of a broodstock reserve where animals could remain undisturbed and able to provide larvae to the Solent system (within and without the extensive European Marine Site (EMS) in the Solent) throughout times of poor recruitment in the wider fished population would provide much greater resilience to collapse. The protection of a broodstock reserve for conservation purposes, is within the remit of the Southern Inshore Fisheries and Conservation Authority (SIFCA) who manage the Solent’s biological resources. The model makes it clear that larvae from any location in the Solent will be widely distributed during its pelagic phase. While this leads to a wide dispersion, or benign re-invasion of this native species, it also means that most locations will not receive the level of spat fall that presumably was normal during periods of much higher abundance. High levels of dispersion work against maintenance of locally high abundance, except for areas where either the combination of hydrodynamics and behaviour concentrate spat at abnormally high concentrations, or where areas are sheltered which limits dispersion. As such, the results support and explain the mismatch between high relative areal extent and low biomass which might have contributed so fundamentally to the over exploitation and decline of native oyster populations world wide (Ermgasson *et al.* 2012). Thus, counter-intuitively perhaps, it may make sense to protect Browndown Bank or Ryde Middle, currently with relatively high natural abundance, rather than exploit these area. Building locally high biomass in these areas may establish natural biogenic reefs (Kennedy & Roberts 1999) and associated diversity which often occur in areas of high exposure (wave and tidal), where they benefit from natural food aggregation (throughput) and offer greater natural resilience to extreme environmental events. There would be an added benefit of production and release of larvae from these locations which would both disperse to the wider area and return in higher relative numbers to maintain a relatively high local population. Thus protection of these high biomass areas may be simultaneously self-sustaining and more productive for the wider area. Warners might also be a good contrasting area to monitor the lower bounds of abundance and dispersion and be a sentinel of range collapse. While an unusual ratio of male v. female phase oysters has been observed in the Solent recently (Kamphausen *et al.* 2011), no disturbances of the reproductive processes have been found in the population (Kamphausen 2012), making it likely that recruitment to the population is limited by larval supply. The approach of establishing broodstock reserves has also been recommended by Laing *et al.* (2005) in a feasibility study of native oyster regeneration in the UK. This study suggests that by managing oyster broodstock areas the SIFCA, could provide long term conservation and fishery benefits, a practical validation of the model’s principles.

### Resilience

The fact that the average larva can cover over 300 km during the pelagic phase, when it is particularly dependent on quick growth, indicates that deleterious conditions in any part of the Solent could impact any or all of the larval population. This would perhaps be most relevant to overall temperature, sea level, food availability, or other natural factors related to climate change such as more frequent wave or storm events. If poor conditions were to prevail in any area of the Solent they would impact the entire larval population, and it would not be possible to use settlement distribution to infer the source of perturbation. It would be useful to contrast the health and size of larvae from sheltered areas such as the Beaulieu Estuary, which had less pelagic exposure, with those from more exposed areas to potentially determine the nature of any systemic changes and the benefits or drawbacks of pelagic exposure. Furthermore, after the pelagic swimming phase, many settlement journeys are long, of the order of several days and may span the model area, often initially outside the Solent. During this time the larvae contact and sample the bed. Therefore, any major bed perturbation, such as dredging, at any point in the Solent during this period may impact spat fall across the entire area. The impacts on larval settlement of dredging in particular locations and other spatially resolved water quality or physical bed disturbance would be good targets for future models of the type proposed in this study.

### Range collapse and refugia

The model results indicate that the perceived shift of the current population from the West Solent into the East Solent is unlikely to be a result of localized deterioration of environmental conditions in the West Solent. Rather, the hydrography appears to be such that those areas where larvae are concentrated most, just happen to lie in the East Solent. These areas would therefore still receive some settlement when overall larval availability has decreased to a point where most areas are no longer supplied, creating the impression of a geographic shift in the population while really being an effect of an overall decrease in larval availability – range collapse. Larval surveys conducted in the Solent in 2010 found that *Ostrea edulis* larvae were present (Kamphausen 2012), but at lower concentrations than during a previous survey in 1984 – 87 (Key 1987). Differences in sampling and analytical techniques mean this is not statistically comparable, but it does follow logic whereby a substantially diminished adult population in 2010 would supply fewer larvae than the population just past peak abundance in 1984.

### Future studies

Our assumptions about bounce settlement behaviour are supported by the literature (Cragg & Gruffyd 1975, Waller 1981, Key 1987) (albeit older studies), but, although difficult, it may be possible to construct an experimental apparatus to observe larval behaviour in the field, effectively by placing the laboratory based apparatus in the earlier studies (an open ended tube) on the bed of the Solent and using a modern underwater camera and appropriate lighting to record the results. Bounce swimming behaviour in increasing pressure, and bed attachment in decreasing pressure would confirm our assumptions and validate our modelling predictions. This experimental set up could also falsify the core hypothesis used in our settlement behavioural model. The impacts of storms and high wave conditions, although usually intermittent, could also be assessed.

Future models would be greatly improved by a joined up set of high resolution water models. In our study, particles often interacted with boundaries and had to be removed from the model. The striking thing is often just how far particles move in these complex tidal environments. The large embayments, along this coast, cause circulatory systems to be set up at each tide – since the larvae are hyper-dispersive (because they swim) they can end up being transported in either direction at each tide. This study supports earlier studies suggesting a linkage between Poole Harbour and the Solent for bivalve larvae (Herbert *et al.* 2012) and the likely link between Solent and the now almost non-existent oyster population in Poole Bay. A fully joined up (and synchronised) water model could be used to map the extent of long range larval transport and improve predictions of population survival and invasions.

## Acknowledgements

We thank the Environment Agency for allowing us to use their hydrodynamic models for research. LK would like to thank the Southern Inshore Fisheries and Conservation Authority and the UK Natural Environmental Research Council for support with the project and doctoral training grant NE/G524160/1

## Attribution

We all contributed equally to the writing, ideas and concepts in the paper. JW and LK lead the writing. JW wrote the code for the behaviour and Lagrangian models and EJ was responsible for the hydrodynamic model.

